# DHA Increases *Angptl4* Gene Expression and Reduces LPL Activity in a PPARγ-dependent manner in Adipocytes

**DOI:** 10.1101/2025.11.15.685896

**Authors:** Patrick V McTavish, Alexander Rajna, Liam H Brown, Reyad Elzaanoun, David M Mutch

**Author notes:** Corresponding Author: David M Mutch, University of Guelph, Department of Human Health Sciences, 50 Stone Road East, Guelph, ON N1G 2W1, Canada.

## Abstract

Omega-3 polyunsaturated fatty acids (N-3 PUFA), specifically eicosapentaenoic acid (EPA) and docosahexaenoic acid (DHA), are well recognized for their triacylglycerol (TAG)-lowering properties. These effects are generally attributed to reduced hepatic lipogenesis and increased β-oxidation; however, the contribution of white adipose tissue (WAT) towards the hypotriglyceridemic properties of N-3 PUFA is less defined. Lipoprotein lipase (LPL) regulates TAG hydrolysis to influence fatty acid uptake into WAT, a process that can be inhibited by angiopoietin-like 4 (ANGPTL4). When re-examining a previous mouse study, we found that mice consuming a diet rich in EPA/DHA had increased WAT *Angptl4* expression in the fasted state compared to a control diet. Therefore, the goal of this study was to explore the role of N-3 PUFA on the regulation of *Angptl4* expression and LPL activity in mouse adipocytes. 3T3-L1 adipocytes treated with DHA (100μM), but not ALA or EPA, increased *Angptl4* expression and reduced LPL activity similar to that observed with a PPARγ agonist (pioglitazone). When *Pparγ* expression was knocked down with siRNA, the ability of DHA and pioglitazone to induce *Angptl4* expression was ablated. Further, DHA- and pioglitazone-induced reductions in LPL activity were mitigated when *Angptl4* expression was silenced. Taken together, these results suggest that DHA regulates LPL activity by increasing *Angptl4* expression in a PPARγ-dependent manner. Our results have uncovered a novel mechanism by which DHA regulates ANGPTL4 to influence LPL-mediated hydrolysis of circulating TAG in adipocytes. Future *in-vivo* studies are necessary to determine the relevance of DHA regulation of ANGPTL4 towards whole-body lipid homeostasis and cardiometabolic health.

**Graphical Abstract:** 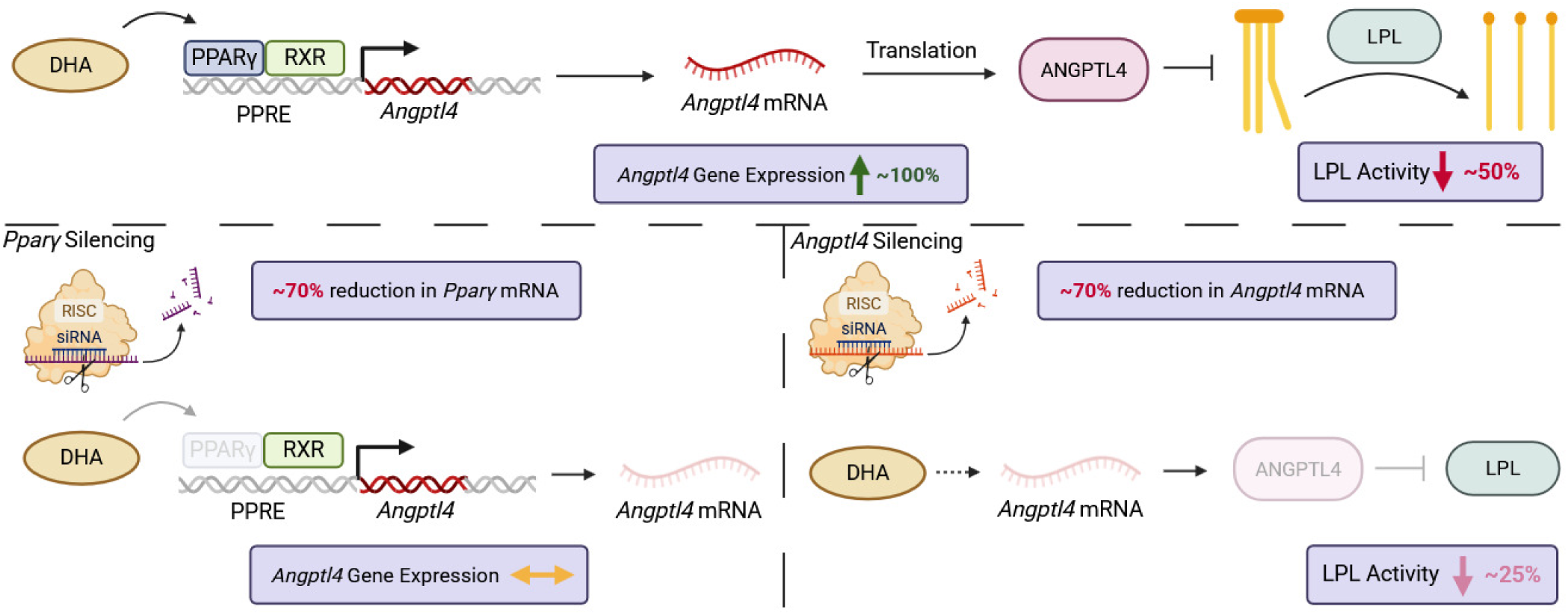

**Highlights:**

- DHA increases *Angptl4* gene expression and reduces LPL activity in 3T3-L1 adipocytes
- Changes in FOXO1 do not explain DHA-induced changes in *Angptl4* gene expression
- Silencing *Pparγ* ablates DHA-induced increases in *Angptl4* gene expression
- Silencing *Angptl4* partially attenuates DHA-induced reductions in LPL activity

## 1. Introduction

Omega-3 polyunsaturated fatty acids (N-3 PUFA), including eicosapentaenoic acid (EPA; 20:5 n-3) and docosahexaenoic acid (DHA; 22:6 n-3), are well known for their triacylglycerol (TAG)-lowering properties [1–4]. This is highly relevant, as circulating TAGs are well-recognized as an independent risk factor for cardiovascular disease (CVD) [5–9], and N-3 PUFA have previously been shown to reduce CVD risk [10–12]. The TAG-lowering properties of N-3 PUFA are generally attributed to reduced *de novo* lipogenesis (DNL) and increased β-oxidation in the liver [2]; however, it remains unclear whether the TAG-lowering effects are also mediated in part through regulation of white adipose tissue (WAT) lipid uptake and/or storage. This is notable because our team recently reported that mice fed a high-fat diet containing EPA/DHA had reduced WAT lipid content compared to control diets [13].

WAT plays a critical role in whole-body lipid homeostasis through the coordinated uptake, storage, and release of fatty acids. Fatty acid uptake into WAT is dependent on lipoprotein lipase (LPL)-mediated hydrolysis of circulating TAGs [14]. Therefore, changes in LPL activity may have significant effects of WAT and whole-body lipid metabolism. We recently collated published literature that, together, suggests that N-3 PUFA can modify LPL activity and alter WAT lipid uptake, although the exact mechanism(s) behind this effect remains unclear [15]. Therefore, we hypothesized that N-3 PUFA regulation of WAT lipid uptake through LPL may provide an additional mechanism underlying the hypotriglyceridemic properties of these beneficial fatty acids.

LPL activity is controlled through both intracellular and extracellular mechanisms. One mechanism involves the angiopoietin-like (ANGPTL) family of proteins, of which ANGPTL3, ANGPTL4, and ANGPTL8 have important roles [16, 17]. ANGPTL proteins function in a coordinated manner to reduce WAT LPL activity in the fasted state and increase it in the fed state [16, 17]. ANGPTL4 is a secreted protein, highly expressed in the liver and WAT [16], that has been associated with numerous outcomes including obesity, inflammation, and abnormal glucose metabolism [18–21]. The primary function of ANGPTL4 is to inhibit LPL post-translationally and reduce its hydrolysis activity [22–24]. The fed-fasted regulation of *Angptl4* expression is partially controlled by insulin signaling, whereby unphosphorylated FOXO1 in combination with glucocorticoid signaling induces *Angptl4* expression in fasted conditions [25]. Previous literature, although not directly examining WAT, supports the idea that N-3 PUFA may modify *Angptl4* expression. Specifically, it was reported that α-linolenic acid (ALA) increased murine cardiac muscle *Angptl4* gene expression, and that DHA increased rat hepatoma cell *Angptl4* expression [26, 27]. Therefore, the goal of this study was to investigate the mechanism by which N-3 PUFA regulate *Angptl4* expression in mouse adipocytes. We hypothesized that this regulation was due to either modifications in insulin signaling, given the fed-fasted regulation of ANGPTL4 [25], or by activation of PPARγ, given that ANGPTL4 is a PPARγ target gene [28, 29]. Lastly, we aimed to connect changes in *Angptl4* expression with functional changes in LPL activity.

## 2. Materials and methods

### 2.1 Brief overview of a previously conducted mouse feeding study

We began the present investigation by revisiting a previous fed-fasting mouse study conducted in our lab [30]. For the purposes of the present study, we only considered mice fed the control (lard) and fish oil diets. Briefly, at 3 weeks of age, male C57BL/6J mice were weaned onto one of two modified AIN93G diets (Research Diets) for 8 weeks. Mice were given *ad-libitum* access to isocaloric low-fat diets (16% kcal from fat) containing either 6.625% lard w/w (lard-diet; 1.5% ALA, 0% EPA, 0% DHA) or 7% menhaden oil w/w (fish-diet; 1.7% ALA, ∼16% EPA, ∼11.5% DHA). The lard diet was supplemented with 0.375% ARASCO oil w/w to match the amount of arachidonic acid naturally present in the fish-diet. Ten mice per diet group had access to food right up to euthanization (fed mice), while another ten mice per diet group were deprived of food for 24h (fasted mice), which began at the start of the light cycle. All mice were euthanized at the beginning of a light cycle. Mice were euthanized by decapitation, and inguinal WAT (iWAT) was collected. Animal protocols were approved by the University of Guelph Animal Care Committee in accordance with the requirements of the Canadian Council of Animal Care.

### 2.2 Cell culture

3T3-L1 preadipocytes were obtained from the American Type Culture Collection (ATCC). Adipocytes were cultured at 37°C, 5% CO_2_, and 100% humidity. After routine passaging, cells were seeded in 6-well plates at a density of 6.0×10^4^ cells per well. Pre-adipocytes were cultured in a basic medium (basic-DMEM) containing Dulbecco’s Modified Eagle Medium (DMEM; CAT # 11965092, Thermo Fisher Scientific) supplemented with 1% penicillin-streptomycin (P/S; CAT# 1037806, Gibco) and 5% heat inactivated fetal bovine serum (FBS; CAT# SH3039602, Cytiva). Media was carefully aspirated and replaced every 2 days. Cells were allowed to grow to 100% confluence, after which confluence was maintained for 48h before induction of differentiation. On the first day of differentiation (i.e., day 0), basic DMEM was replaced with differentiation media, which consisted of basic-DMEM supplemented with 1μM dexamethasone (DEX; CAT# D4902, Sigma), 0.5mM 3-isobutyl-1-methylxanthine (IBMX; CAT# I5879, Sigma), and 1µg/mL of insulin (CAT# I9278, Sigma). On day 2 post-induction of differentiation (PID), differentiation media was removed and a maintenance media consisting of basic-DMEM supplemented with 1µg/mL insulin was added. Maintenance media was replaced on day 4 PID and was then changed to basic-DMEM from day 5 PID to day 8 PID. Fatty acid treatments were prepared by adding 2% w/v fatty acid-free BSA (CAT# A3803, Sigma) to basic-DMEM, and then diluting a fatty acid stock solution prepared in 100% EtOH in the media to a final concentration of 100μM. Pioglitazone (PIO) treatment was prepared by diluting a PIO stock prepared in 100% DMSO in the media to a final concentration of 1μM. The control condition media was supplemented with fatty acid-free BSA. All treatment, including control, were matched to 1:100 EtOH and 1:1000 DMSO. All cell studies were conducted in different passages and lots to ensure results were generalizable. Using these general methods for culturing and differentiating 3T3-L1 preadipocytes, we subsequently conducted the following series of experiments:

#### 2.2.1 Experiment 1 – Fatty Acid Treatments

Differentiated 3T3-L1 adipocytes were treated with 100µM of the following fatty acids: palmitic acid (PA; 16:0), α-linolenic acid (ALA; 18:3n-3), eicosapentaenoic acid (EPA; 20:5n-3), or docosahexaenoic acid (DHA; 22:6n-3) for 48 hours beginning on day 8 PID.

#### 2.2.2 Experiment 2 – Fatty Acid and PIO Treatments

Treatment groups in this experiment were similar to Experiment 1, with the addition of 1µM pioglitazone (PIO; CAT# E6910, Sigma), resulting in a total of 6 treatment groups.

#### 2.2.3 Experiment 3 – siRNA Treatments

Differentiated 3T3-L1 adipocytes were treated with siRNA targeting *Pparγ* or *Angptl4,* as described in Section 2.7. The treatment groups were reduced to include only the control condition, DHA, and PIO as described in Experiment 2, resulting in a total of 6 treatment groups in a 2×3 design.

### 2.3 RNA extraction, quantification, and RT-qPCR

Media was aspirated and the cells were washed twice with 1× PBS. Total RNA was extracted from cells using the QIAGEN RNeasy Mini Kit^TM^ (CAT# 74106, Qiagen), according to the manufacturer’s protocol. RNA concentration and 260/230 absorbance ratio was measured with a Thermo Scientific NanoDrop 2000 spectrophotometer. RNA extraction and quantification for mouse WAT samples was performed in a similar manner to that of cultured adipocytes, and is explained in more detail in [30].

Extracted RNA was reverse transcribed into cDNA using the Applied Biosystems High-Capacity cDNA Reverse Transcription Kit (CAT# 4374966, Applied Biosystems). according to the manufacturer’s protocol. The following cycling conditions were used: one denaturing cycle at 95°C for 30 s, followed by 40 cycles of 95°C for 4 s and 55.9°C for 4 s. *Nono* was used as the housekeeping reference gene. Data was analyzed using the ΔΔCt method. Gene expression data is represented as fold changes relative to the control condition. The primers used to measure *Nono, Angptl4, Pparγ, Lpl, Atgl* and *Hsl* expression are provided in **Supplemental Table 1.**

### 2.4 Protein extraction, quantification, and western blot

Protein was extracted from cells using RIPA lysis buffer (CAT# 0089900, Thermo Fisher Scientific) supplemented with a 1:100 Halt™ Protease and Phosphatase Inhibitor Cocktail 100× (CAT #: 1861281, Thermo Fisher Scientific). Cell lysates were then centrifuged at 10,000rcf for 10 minutes, the supernatant was collected, and cellular debris was discarded. Protein yields were quantified using the BCA assay according to the manufacturer’s protocols (CAT# 23209, Thermo Fisher Scientific). Sample absorbance was measured at 562nm using a Spectramax M2e microplate reader, and protein concentrations were used to prepare 1mg/mL denatured protein samples in 4× lammeli sample buffer (CAT# 1610747, Bio-Rad) supplemented with β-mercaptoethanol according to the manufacturer’s instructions.

Equivalent amounts of denatured cell lysate (20 μg) were loaded and separated by SDS-PAGE before transferring to a nitrocellulose membrane (CAT# 1620115, Bio-Rad). Prior to blocking or probing with a primary antibody, membranes were stained with a Ponceau-S solution (CAT# P7170, Sigma) and images were taken. Membranes were then blocked in 5.0% BSA in tris-buffered saline containing 0.1% Tween 20 (TBST), followed by an overnight incubation at 4°C with primary antibody diluted in 5.0% BSA/TBST. Subsequently, membranes were washed in TBST and incubated with a secondary antibody diluted in 5.0% BSA/TBST for 2 hours. Protein bands were visualized with Pierce ECL western blotting substrate (CAT# 32106, Thermo Fisher Scientific) in a FluorChem HD2 Imaging System. The following primary antibodies were used: α-tubulin (CAT#: ab7291, Abcam, 1:1000 dilution), total AKT (CAT# 9272S, Cell Signaling, 1:1000 dilution), phosphor-Ser^473^ AKT (CAT# 9271S, Cell Signaling, 1:1000 dilution), total FOXO1 (CAT# 9454S, Cell Signaling, 1:1000 dilution), phospho-Ser^256^ FOXO1 (CAT# 9461S, Cell Signaling, 1:1000 dilution), and LPL (CAT# AF7197, biotechne, 1:1000 dilution). The secondary antibodies used were Goat Anti-Rabbit IgG HRP conjugate (CAT# 1706515, Bio-Rad Laboratories, 1:3000 dilution), Goat Anti-Mouse IgG HRP conjugate (CAT# 1706516, Bio-Rad Laboratories, 1:3000 dilution), and Mouse Anti-Goat IgG HRP conjugate (CAT# HAF017, biotechne, 1:1000 dilution). Protein band intensities were quantified by AlphaView SA software (Cell Biosciences Inc).

### 2.5 Lipoprotein lipase activity assay

LPL activity in 3T3-L1 adipocytes was assayed using the Abcam fluorometric lipoprotein lipase activity assay (CAT# ab204721, Abcam), according to the manufacturer’s protocol. LPL activity was measured using both the whole cell fraction (cell lysate; as per the manufacturer’s protocol) and the heparin fraction using a protocol adapted from Makoveichuk et al [31]. For the heparin fraction, differentiated adipocytes were washed twice with cold 1× PBS and subsequently incubated with a 100IU/mL solution of heparin (CAT# 375095, Sigma) in DMEM containing 0.2% fatty acid free BSA (CAT# A3803, Sigma) for 30 minutes at 4°C. Fluorescence was measured using a Spectramax M2e microplate reader set to kinetic mode, measuring Ex/Em 482/515 nm for 2 hours at 37°C. Data for this assay is represented as the rate of enzyme activity (mU), which corresponds to the amount of LPL that generates 1.0 nmol of fatty acid product per minute at pH 7.4 and 37°C. We further normalized this data to total protein content from cell lysates.

### 2.6 Oil red O lipid staining

Neutral lipids in adipocytes were stained with Oil-Red O. Briefly, a working solution of Oil-Red O was prepared by combining 9 parts of an Oil-Red O stock (CAT# O0625, Sigma) with 6 parts dH_2_O. Media was aspirated from cells, and the cells were then washed twice with 1× PBS. 2mL of 10% formalin in PBS (CAT# SF100-4, Thermo Fisher Scientific) was added to each well and allowed to incubate for 5 minutes before being removed. Then, 2mL of formalin was added again and allowed to incubate for at least one hour. Formalin was then removed and cells washed once with a 60% isopropanol solution and allowed to dry completely. From there, 1mL of the Oil-Red O working solution was added to each well and allowed to incubate for 10 minutes. The Oil-Red O solution was then aspirated, and wells were washed 4× with 2mL of dH_2_O. Images were taken using an AMG-EVOS XL microscope set to 10× magnification.

### 2.7 Reverse Transfection siRNA

Reverse transfection siRNA gene silencing was performed using the Silencer Select siRNA system from Life Technologies, according to the manufacturer’s protocol. Our protocol was based on the protocol described by Isidor et al [32], with minor modifications. On day 8 PID, differentiated adipocytes grown in 6-well plates were dissociated from the wells using a trypsin-EDTA solution (CAT# 25200056, Gibco). From there, cells were seeded at a density of 2.0×10^5^ cells/well in a 48-well plate precoated with gelatin (CAT# G1393, Sigma), and a mixture of siRNA (90nM) and Lipofectamine RNAiMAX (CAT# 13778030, Life Technologies) prepared in Opti-MEM (CAT# 31985062, Gibco). The specific siRNAs used were PPARy (CAT# 4390771, siRNA ID s72013, Life Technologies), ANGPTL4 (CAT# 4390771, siRNA ID s81455, Life Technologies), and the Silencer Select negative control siRNA #1 (CAT# 4390843, Life Technologies). Cells were pre-treated with siRNA for 48 hours (i.e., until day 10 PID), at which point the fatty acid or PIO treatments were added for a further 48 hours (i.e., until day 12 PID).

### 2.8 Cytotoxicity testing

Cytotoxicity of treatments on 3T3-L1 adipocytes was tested using a lactate dehydrogenase assay, in accordance with the manufacturer’s protocol (CAT# G1780, Promega). Cell media samples were diluted 5-fold and assayed in duplicate. Unconditioned basic-DMEM was used as a blank to control for any LDH which may be naturally present in culture media.

### 2.9 Statistical analysis

Data was analysed using a repeat measures one-way or two-way ANOVA to assess main effects and interaction effects. When appropriate, Tukey’s HSD multiple comparison tests were carried out to assess pairwise comparisons. Statistical significance was determined at p <0.05. Outliers were identified using the ROUT test. When an outlier was identified, the entire passage of treatments was removed from analysis to maintain the requirement of equal group sizes for the repeat-measures ANOVA. All analyses and graphing were conducted using Prism Graphpad 10.

## 3. Results

### 3.1 A diet rich in EPA/DHA increases iWAT Angptl4 gene expression

We first assessed whether the consumption of a diet containing fish oil (rich in EPA/DHA) affected *Angptl4* gene expression compared to a lard diet containing no EPA or DHA in fed and fasted male C57BL/6J mice. Using tissue samples collected from a previous mouse feeding study [30], we found a strong trend for an interaction (p = 0.06) between diet and nutritional state on iWAT *Angptl4* gene expression, as well as highly significant main effects for diet (lard vs. fish oil; p < 0.01) and nutritional state (fed vs. fasted; p <0.01) on iWAT *Angptl4* gene expression. This suggested that a diet containing EPA/DHA had a stronger effect on *Angptl4* expression in the fasted state than a diet lacking these N-3 PUFA. Although the interaction did not meet our p<0.05 cut-off, we conducted post hoc tests due to the strong trend and found that mice fed the fish oil diet had a significantly higher expression of *Angptl4* relative to mice fed the lard diet in the fasted state (p <0.01), with no difference in *Angptl4* expression in fed mice (**Figure 1**).

**Figure 1:**
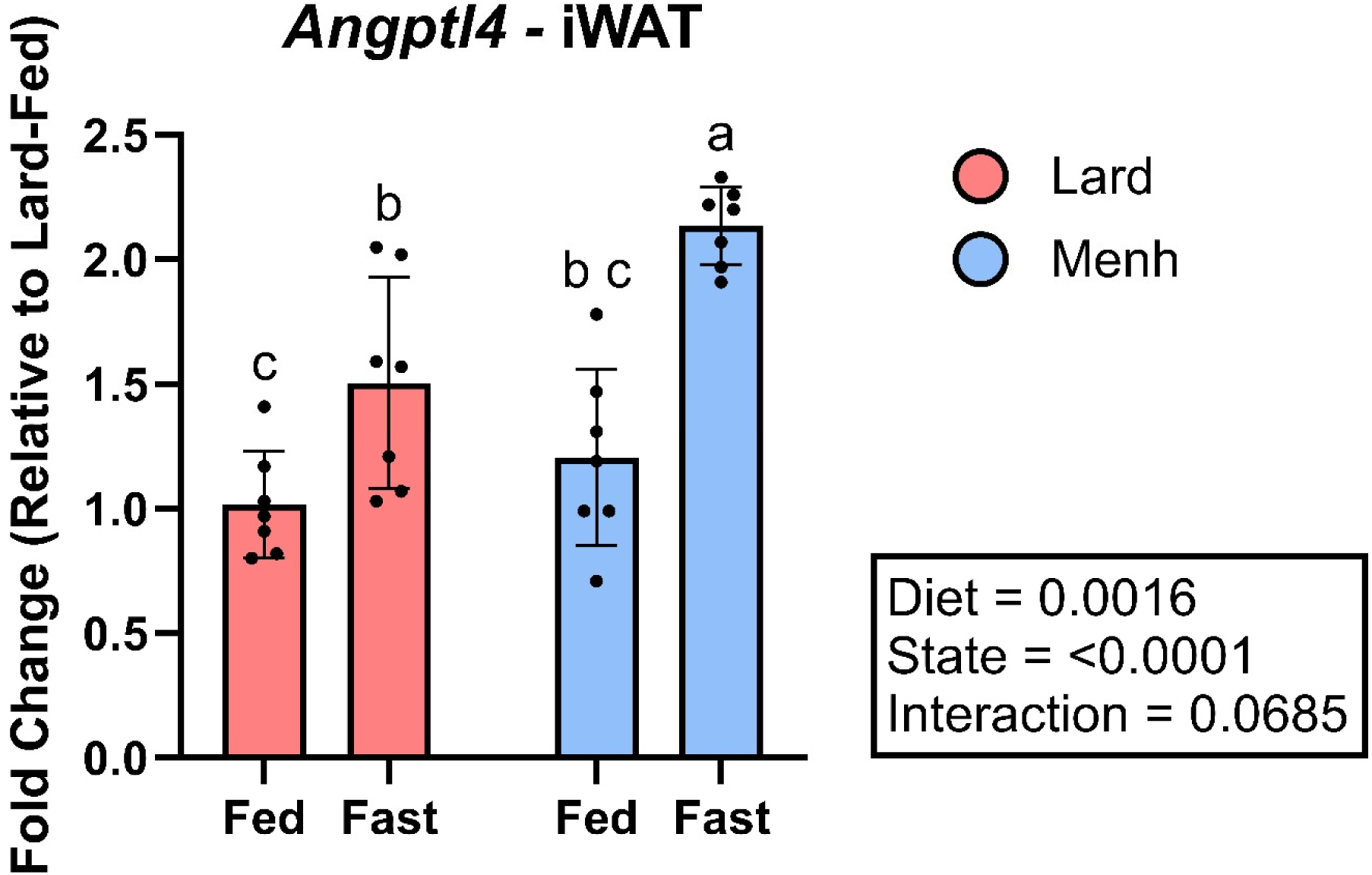
*Angptl4* expression in iWAT samples from mice fed a lard or fish oil diet for 8 weeks. Gene expression data was normalized to *Nono,* and fold changes were calculated relative to the lard-fed group. Data are presented as mean ± SD (n=7 mice per group). *Angptl4* expression was analyzed by two-way ANOVA, followed by a Tukey’s post hoc test.

### 3.2 DHA increases Angptl4 gene expression in 3T3-L1 Adipocytes independent of changes in FOXO1

All subsequent investigations into the relationship between N-3 PUFA and *Angptl4* expression were conducted in 3T3-L1 mouse preadipocytes. After 48h of treatment, DHA resulted in a ∼2-fold increase in *Angptl4* gene expression in differentiated adipocytes relative to control cells (p <0.01; **Figure 2**). No changes in *Angptl4* expression were observed with PA or ALA, while EPA showed an intermediate yet statistically insignificant increase.

**Figure 2:**
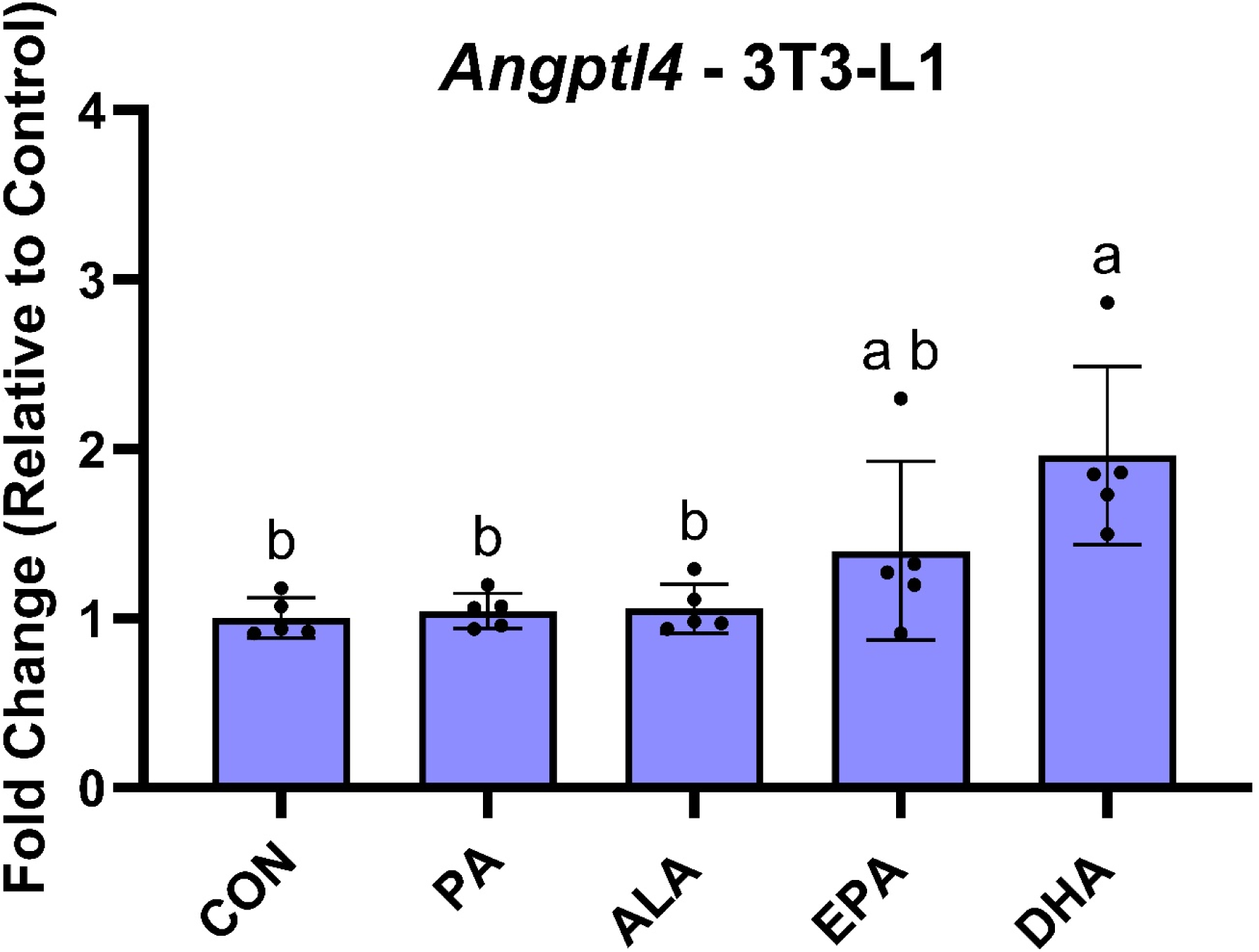
*Angptl4* gene expression in differentiated 3T3-L1 adipocytes treated with different fatty acids. Gene expression data was normalized to *Nono*, and fold changes were calculated relative to the control (CON) group. Data are presented as mean ± SD (n = 5 passages of cells). *Angptl4* expression was analyzed by a repeat measures one-way ANOVA, followed by a Tukey’s post hoc test.

Due to the known regulation of *Angptl4* gene expression by FOXO1, we next investigated the effects of DHA on total and phospho-Ser^256^ levels of FOXO1 in both basal and insulin-stimulated conditions. Neither total nor phospho-Ser^256^ levels of FOXO1 were altered in the basal state by the different fatty acids (**Figures 3A-B**). Insulin stimulation increased phospho-Ser^256^ levels of FOXO1 in all conditions, with a 1-way ANOVA revealing a significant effect of insulin on the ratio of pFOXO1 Ser^256^ / total FOXO1 (p=0.03) as well as pAKTSer^473^ / total AKT (p=0.03). However, subsequent post-hoc analysis revealed that the insulin-stimulated increase in pFOXO1 did not differ between the fatty acid treatments and the control. However, a statistical difference was observed between DHA and PA treatments (Figures 3C-D). Collectively, these results suggest that the DHA-induced increase in *Angptl4* expression was not likely mediated by changes in insulin signaling in adipocytes.

**Figure 3:**
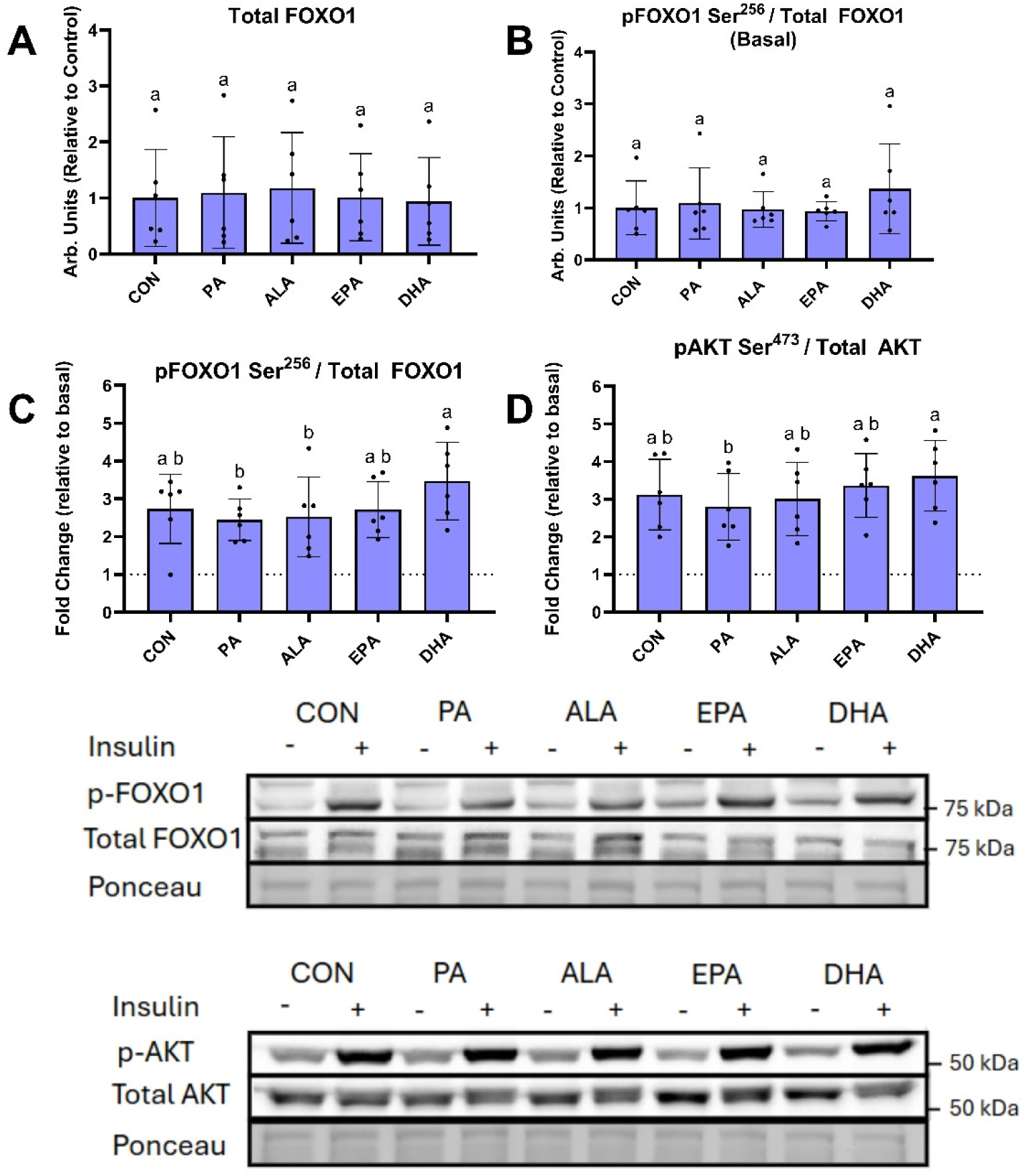
Markers of insulin signaling in differentiated 3T3-L1 adipocytes treated with different fatty acids. (A) The relative abundance of total FOXO1 protein. (B) The relative ratio of pFOXO1 ser^256^ to total FOXO1 under basal (non-insulin stimulated) conditions. (C) The fold-change in the ratio of pFOXO1 to total FOXO1 and (D) The fold-change in the ratio of pAKT ser^473^ to total AKT between basal and insulin stimulated conditions; The dotted line at y=1 represents the phosphorylation level of the basal conditions for each treatment. (E-F) representative blots, (-) indicates basal, (+) indicates insulin stimulated. Data are presented as mean ± SD (n = 6 passages of cells). Data were analyzed by repeated measures one-way ANOVA followed by a Tukey’s post-hoc test.

### 3.3 DHA and pioglitazone-induced increases in Angptl4 expression in 3T3-L1 adipocytes are associated with decreases in cellular LPL activity

We next investigated the functional impact of changes in *Angptl4* expression and the role of PPARγ, since *Angptl4* is a known target of this transcription factor [28]. After 48 hours of treatment with the different fatty acids or pioglitazone (PIO), we did not observe differences in Oil-Red O stain of intracellular lipids (**Figure 4**). Nevertheless, when we reran all fatty acid treatments and included pioglitazone (PIO) as a positive control for PPARγ stimulation, we again observed a significant ∼2-fold increase in *Angptl4* expression in differentiated adipocytes treated with DHA (p <0.01, **Figure 5A**). The magnitude of this increase was similar to that observed in cells treated with PIO (p<0.01). While a small increase was observed with EPA in this second round of experiments, there was still no effect seen with PA or ALA. A post-hoc analysis revealed the increase in *Angptl4* expression was not different between DHA and PIO.

**Figure 4:**
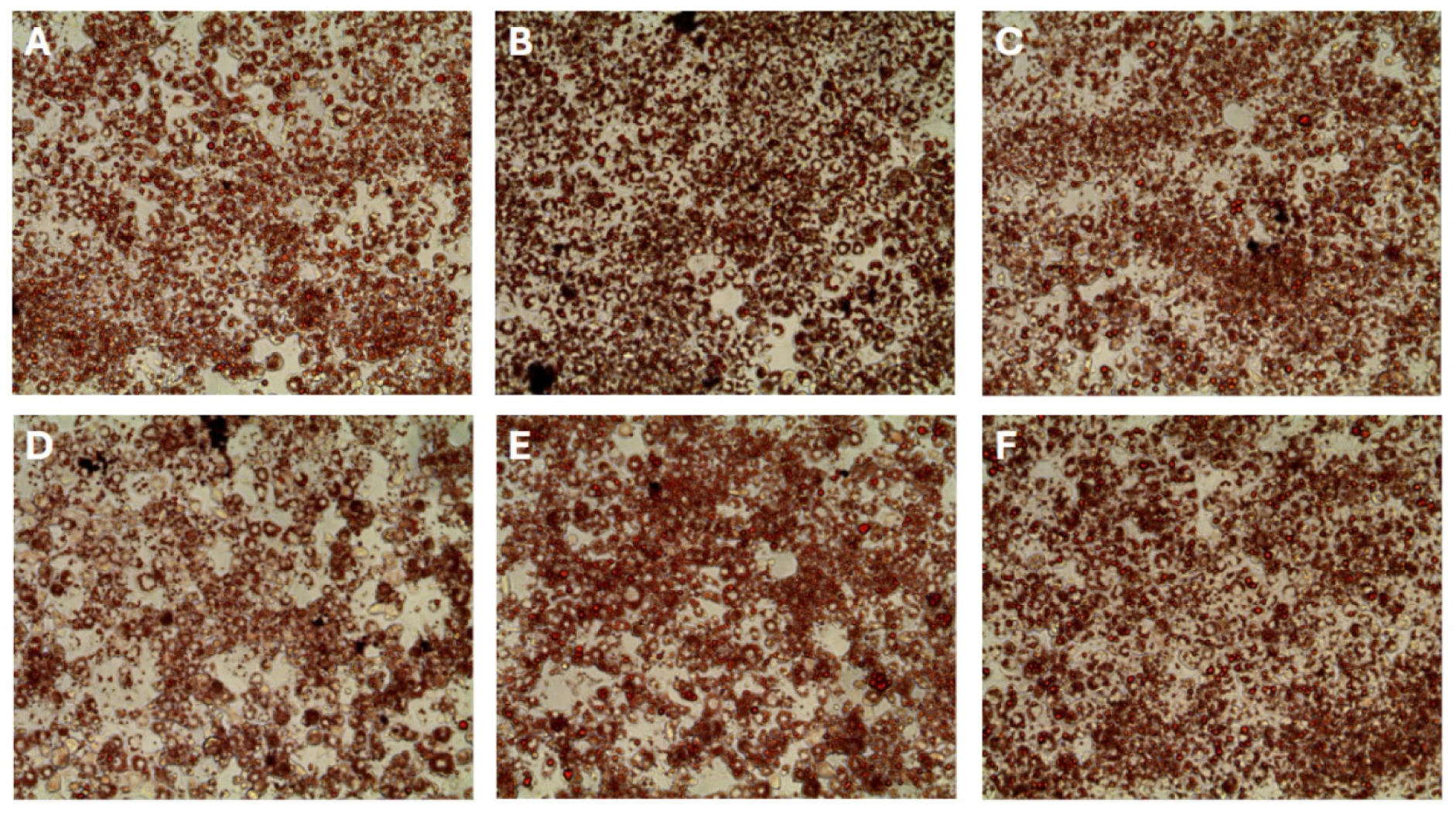
Staining for neutral lipids in differentiated adipocytes treated with different fatty acids. (A) Control, (B) 100µM PA, (C) 100µM ALA, (D) 100µM EPA, (E) 100µ M DHA, and (F) 1µM PIO. Images were taken at 10× magnification.

**Figure 5:**
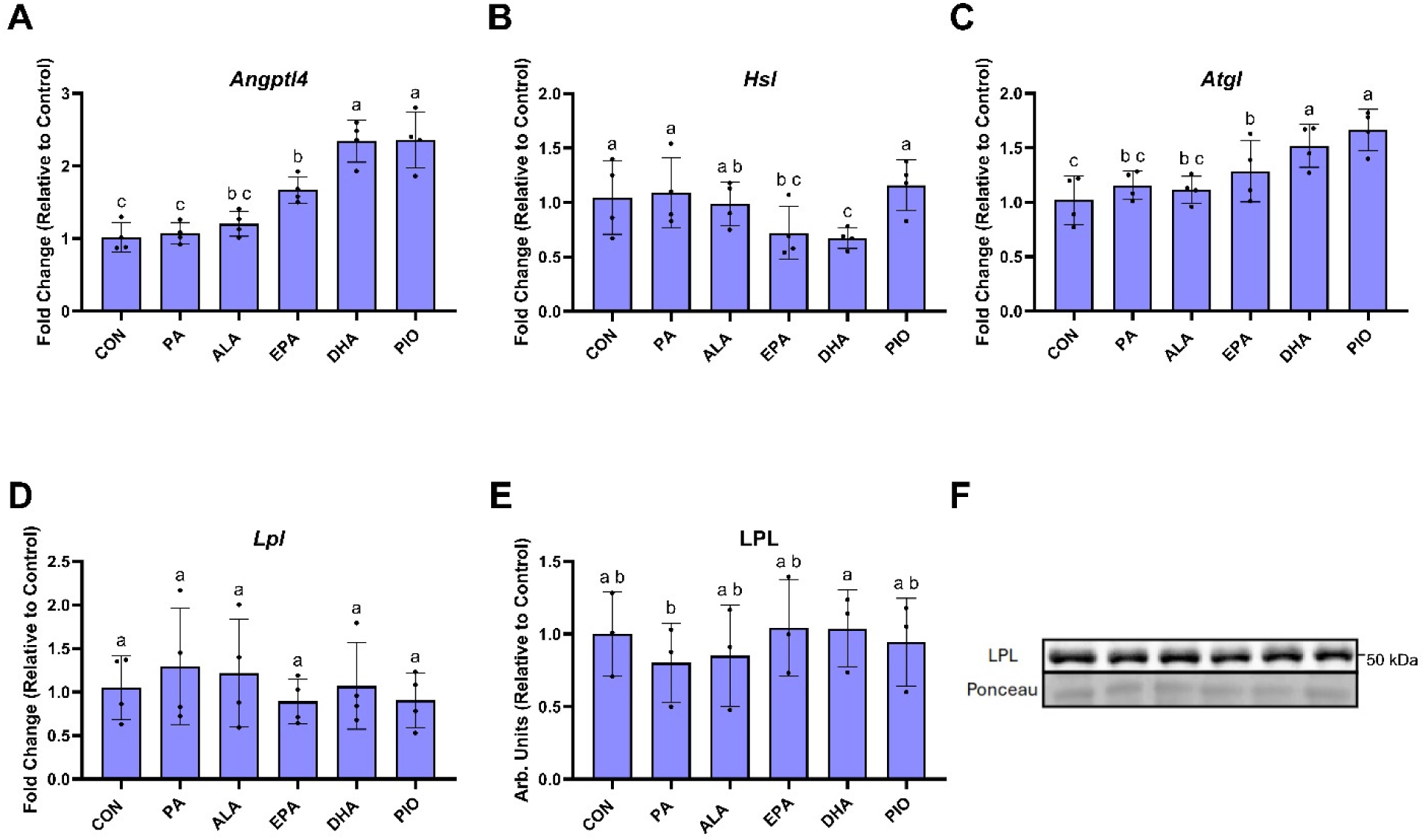
Gene and protein expression in differentiated 3T3-L1 adipocytes treated with fatty acids or PIO. (A-D) *Angptl4, Hsl, Atgl,* and *Lpl* gene expression was normalized to *Nono*, with fold changes calculated relative to CON (n=4). (E-F) Relative abundance of LPL protein was assayed by western blot (n=3). Data are presented as mean ± SD, and all data was analyzed by repeat measures one-way ANOVA followed by a Tukey’s post-hoc test.

Concomitant to these changes in *Angptl4* expression, an approximate ∼50% reduction in protein-normalized cellular LPL activity was observed in adipocytes treated with either DHA (p = 0.01) or PIO (p = 0.02; **Figure 6A**). No effect on LPL activity was observed with PA, ALA or EPA (**Figure 6A**). Similarly, a significant reduction in heparin-releasable LPL activity was observed for cells treated with PIO (P < 0.01), with an intermediate effect observed with DHA. The other fatty acids had no effect on heparin-releasable LPL activity (**Figure 6B**). Although minor changes in gene expression were observed with some cellular lipases (**Figure 5B-D**), these changes did not align with DHA- and PIO-mediated changes in LPL activity. Specifically, we observed a reduction in *Hsl* with EPA (p = 0.04) and DHA (p = 0.01), with no effect with PIO (**Figure 5C**). We also observed an increase in *Atgl* expression with EPA (p = 0.02), DHA (p <0.01), and PIO (p < 0.01) (**Figure 5D**). Finally, no changes were observed in adipocyte *Lpl* gene expression or LPL protein abundance (**Figures 5D-F**).

**Figure 6:**
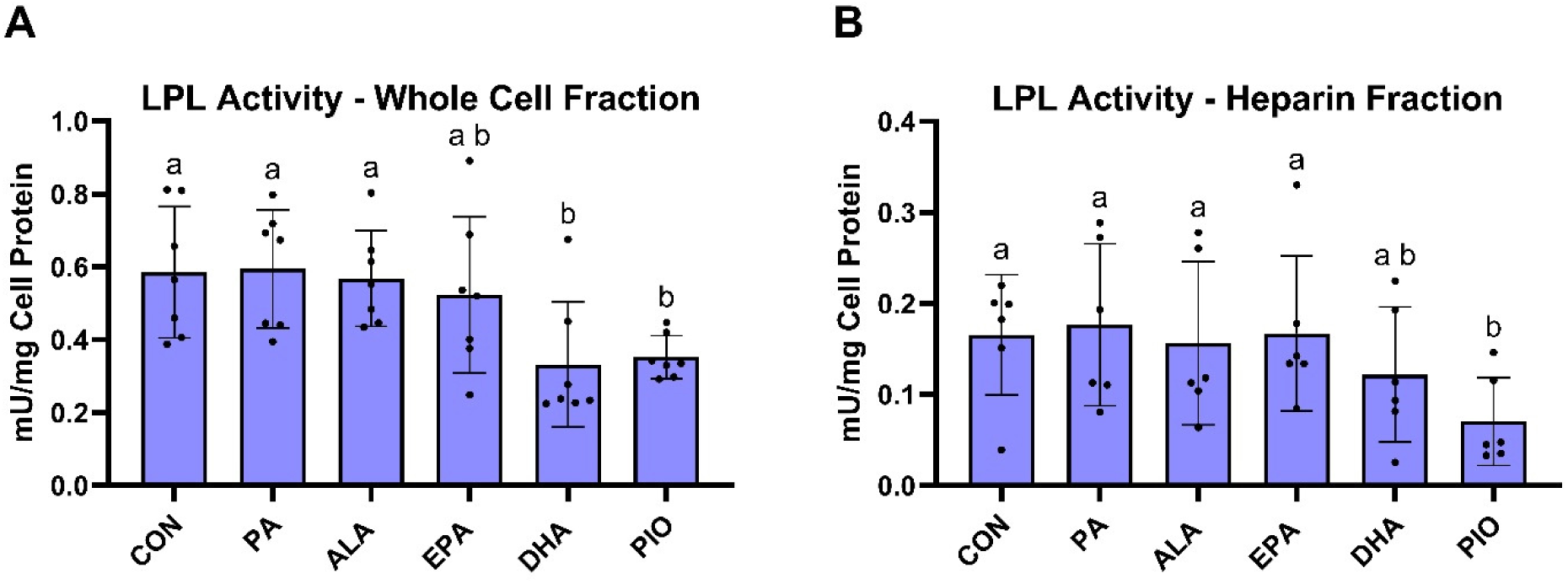
LPL activity in differentiate 3T3-L1 adipocytes treated with fatty acids or PIO. (A) LPL activity measured in whole cell extracts expressed as milliunit (mU) per mg of cell lysate protein (n=7). (B) Heparin-releasable LPL activity expressed as milliunit (mU) per mg of cell lysate protein (n=6). Data are presented as mean ± SD, and all data was analyzed by repeat measures one-way ANOVA followed by a Tukey’s post-hoc test.

### 3.4 PPARy silencing ablates DHA and pioglitazone-induced increases in Angptl4 expression in 3T3-L1 adipocytes

Due to the similar effects observed with DHA and PIO, we next investigated the specific role of PPARγ by knocking down its cellular expression with siRNA. An approximate 70% knockdown in *Pparγ* gene expression was achieved with siRNA (**Figure 7A**). In cells treated with a scramble siRNA, DHA and PIO caused small but significant reductions in *Pparγ* gene expression (**Figure 7A**). However, these treatment effects were ablated in cells treated with siRNA targeting *Pparγ*. When examining *Angptl4* expression in these cells (**Figure 7B**), we observed strong increases in *Angptl4* expression with DHA (p<0.01) and PIO (p<0.01) in the scramble condition (similar to that seen in **Figure 5A**), and a significant difference between DHA and PIO (p = 0.03). However, when *Pparγ* was knocked down, the effects of DHA and PIO on *Angptl4* expression were largely ablated (**Figure 7B**). Additionally, there was no difference in *Angptl4* expression between DHA and PIO when *Pparγ* was silenced (**Figure 7B**). Lastly, we observed a main effect of siRNA (p < 0.01) to reduce *Lpl* gene expression in *Pparγ* silenced cells (**Figure 7C**). Taken together, this suggests that DHA and PIO work, at least in part, through PPARγ to promote *Angptl4* gene expression in differentiated adipocytes.

**Figure 7:**
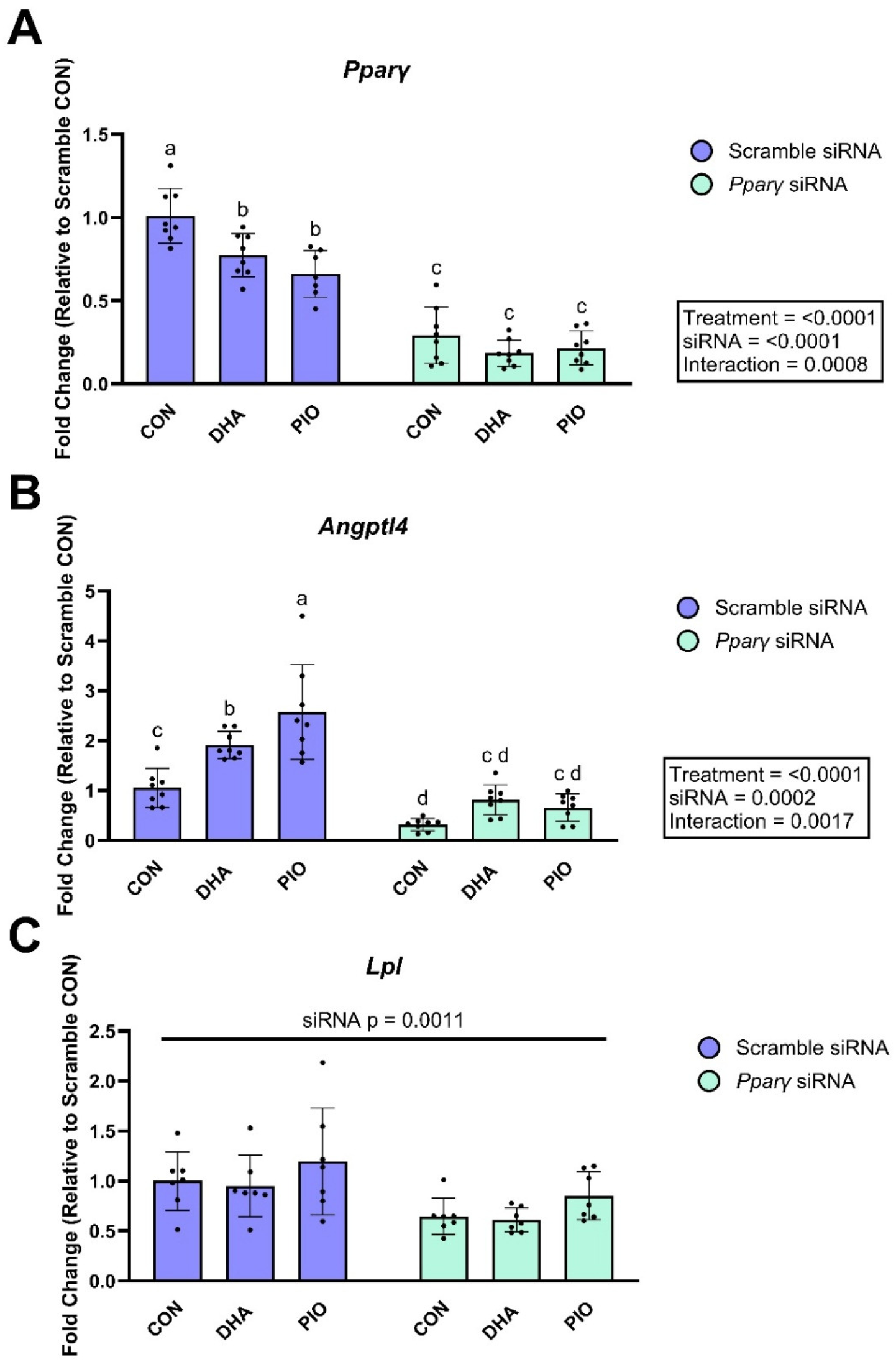
Gene expression data in differentiated 3T3-L1 adipocytes treated with *Ppary* (A) *Pparγ*, (B) *Angptl4*, and (C) *Lpl* gene expression were normalized to *Nono*, with fold changes calculated relative to the Scramble-CON group. All data are presented as mean ± SD (n = 8 passages of cells, one outlier passage removed in (C)). Data was analyzed by repeat-measures two-way ANOVA followed by a Tukey’s post-hoc test.

### 3.5 ANGPTL4 silencing reduces the extent to which DHA and pioglitazone reduce intracellular LPL activity in 3T3-L1 adipocytes

We next examined the role of *Angptl4* on DHA and PIO reductions in LPL activity. Similar to our previous experiments, DHA and PIO increased *Angptl4* expression by ∼2-fold (both p<0.01) when cells were treated with a scramble siRNA (**Figure 8A**). An approximate 70% reduction in *Angptl4* expression was observed in the control treatment when *Angptl4* was silenced with siRNA (**Figure 8A**). Despite this, we observed a borderline significant difference between CON and DHA (p = 0.06) and a significant difference between CON and PIO (p<0.01) in the siRNA group. LPL activity was affected by both treatment (p < 0.01) and siRNA (p < 0.01); however, a significant interaction was not detected (p = 0.99; **Figure 8B**). Both DHA (p = 0.03) and PIO (p = 0.02) reduced LPL activity by ∼50% in the scramble siRNA condition, similar to the results shown in **Figure 6A**. While DHA (p = 0.02) and PIO (p = 0.01) also reduced LPL activity relative to control when *Angptl4* was silenced, the effect size was notably smaller **(Figure 8C)**. When representing the change in LPL activity induced by DHA or PIO relative to the control group (which was set to zero), we observed a significant main effect (p = 0.02) indicating that when *Angptl4* was silenced, the effects of DHA and PIO on LPL activity were minimized compared to the scramble condition **(Figure 8C)**. These effects were independent of changes in *Lpl* gene expression (**Figure 8D**).

**Figure 8:**
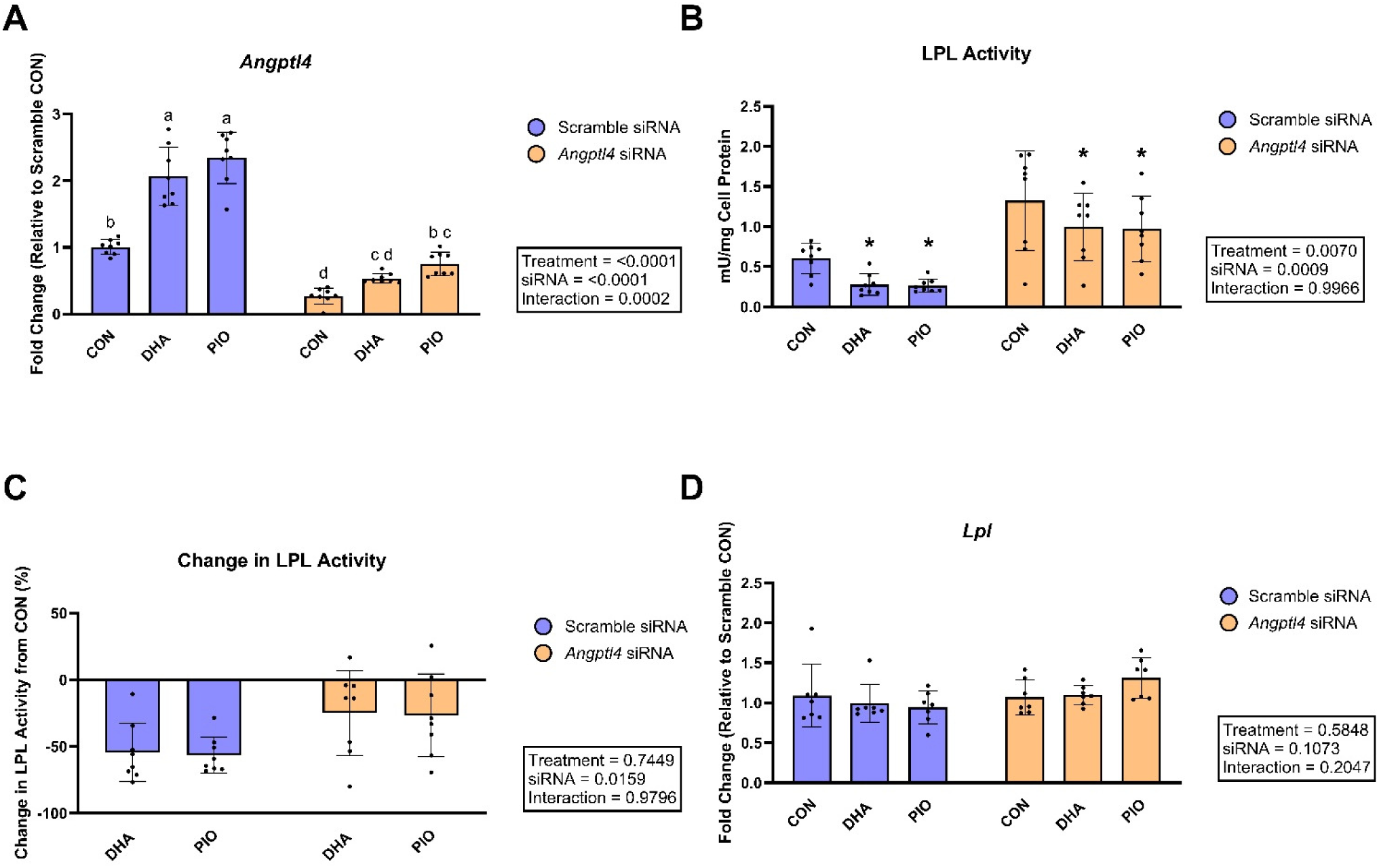
LPL activity and gene expression data in differentiated 3T3-L1 adipocytes treated with *Angptl4* siRNA. (A) *Angptl4* gene expression was normalized to *Nono*, with fold changes calculated relative to the Scramble-CON group. (B) LPL activity was measured in whole cell extracts, expressed as milliunit (mU) per mg of cell lysate protein. (C) Change in LPL activity relative to the CON group for each siRNA condition. (D) *Lpl* gene expression was normalized to *Nono*, with fold changes relative to the Scramble-CON group. All data are presented as mean ± SD (n=8 passages of cells; one outlier passage removed in (D)). Data was analyzed by repeat-measures two-way ANOVA followed by Tukey’s post-hoc test (A), Bonferroni’s post-hoc test (B), and simple main effects (C). Post-hoc significance is denoted as follows: * = p<0.05, ** = p<0.01.

## 4. Discussion

The present investigation used both mice and cultured adipocytes to uncover that N-3 PUFA regulate *Angptl4* expression and LPL activity in adipocytes via PPARγ. Specifically, we found that adipocytes treated with DHA, but not ALA or EPA, had reduced cellular LPL activity in conjunction with increased *Angptl4* gene expression. Neither changes in FOXO1 protein expression nor phosphorylation status were able to explain these findings. In contrast, we found that when *Pparγ* expression was silenced, DHA-mediated increases in *Angptl4* expression were ablated. Similarly, when *Angptl4* was silenced, DHA was unable to reduce LPL activity to the same extent as cells treated with a scramble siRNA. Taken together, this work has uncovered that DHA reduces LPL activity through PPARγ regulation of *Angptl4* expression. Although our findings appear to be opposite to our initial hypothesis that the hypotriglyceridemic properties may be associated with N-3 PUFA up-regulation of LPL activity, we believe that the present findings have uncovered a mechanism by which N-3 PUFA regulate WAT lipid uptake and, potentially, whole-body lipid metabolism. Moreover, these findings provide mechanistic insight into our previous work showing that a diet rich in EPA/DHA reduced TAG content and adipocyte size in visceral AT [13]

Our results align with previous findings in rat hepatoma cells demonstrating that DHA can increase *Angptl4* gene expression [27]. However, our results represent the first study, to our knowledge, reporting that DHA specifically elicits an increase in *Angptl4* expression both in mouse WAT and cultured adipocytes. Furthermore, our results are in agreement with the established role of ANGPTL4 to reduce fasted-state WAT LPL activity [16], since we found that a diet containing fish oil (EPA and DHA rich) increased iWAT *Angptl4* expression in the fasted state and not the fed state. In contrast with our results, a study by Brands et al. [27] demonstrated that the concomitant infusion of fish oil with insulin in humans attenuated the insulin-induced reductions in circulating ANGPTL4. In our mouse study, we did not see an effect of the fish oil diet on *Angptl4* expression in iWAT in the fed state; however, we acknowledge that circulating levels of ANGPTL4 may not reflect *Angptl4* expression in iWAT due to the contribution of other tissues such as the liver [16]. While it remains unclear what portion of circulating ANGPTL4 originates from WAT depots, data from adipose-specific ANGPTL4 KO mice suggests that adipose tissue-derived ANGPTL4 plays an important role in overall metabolic health [33, 34].

*Angptl4* gene expression is known to be regulated by multiple transcription factors, including PPARs and FOXO1 [25, 28]. For the purposes of this study, we focused on FOXO1 and PPARγ, given that DHA in particular has been shown to both modify insulin sensitivity in some contexts [35] and is recognized to be a PPARγ ligand [36]. Our results indicate that DHA does not appear to alter adipocyte insulin signaling pathways, as reflected by the lack of change in total or phosphorylated FOXO1 and AKT. While these findings may appear to contrast with previous reports suggesting that fish oil can improve insulin sensitivity in humans [35], it is possible that 3T3-L1 mouse adipocytes do not fully capture an *in-vivo* model. Regardless, if DHA-induced increases in *Angptl4* expression were mediated by FOXO1, we would have expected to observe either an increase in total FOXO1 (which is transcriptionally active) or a reduction in basal phosphorylation of FOXO1 (which is transcriptionally inactive) [25]. However, neither of these changes were observed in our experiments. An important caveat is that the role of FOXO1 cannot be discounted *in vivo* without further investigation and consideration of chronic versus acute DHA treatment effects. For example, it is possible that N-3 PUFA-mediated increases in insulin signaling could decrease *Angptl4* expression in the fed state via increased FOXO1 phosphorylation [25]. Conversely, the role of FOXO1 in this relationship may be emphasized in the obese state, rather than the healthy state. N-3 PUFA-mediated increases in insulin sensitivity, as seen in the context of T2D [35], may attenuate the elevated levels of ANGPTL4 seen with insulin resistance [37]. Therefore, the relationship between N-3 PUFA and *Angptl4* described herein may be modified in insulin resistant or obese states, representing an avenue for further study.

In contrast, our data indicates DHA regulation of *Angptl4* expression is dependent, at least in part, on PPARγ. This mechanism of action is plausible given that *Angptl4* contains a PPAR-response element in its promoter region [29], and DHA is known to be an endogenous PPARγ ligand. While there is a small visual increase in *Angptl4* expression induced by DHA and PIO in PPARγ-silenced cells, it is important to note that these differences were not statistically significant. We believe that this minor increase can be attributed to the fact that siRNA does not produce a 100% KO in *Pparγ* expression. This is particularly notable given the relatively high expression level of *Pparγ* in adipocytes. As such, it is plausible that any residual *Pparγ* expression in the cells may be sufficient to allow for a small effect on *Angptl4* expression with DHA and PIO. Alternatively, we can’t exclude a possible role for PPAR delta (PPARδ), given that it is also expressed in 3T3-L1 adipocytes and was reported to increase *Angptl4* gene expression [38]. Regardless, the lack of a significant difference between DHA and PIO is further evidence that DHA may be regulating *Angptl4* through PPARγ. Unexpectedly, we also found that DHA (as well as PIO) reduced *Pparγ* gene expression. However, this aligns with a previous study demonstrating that troglitazone reduced *Pparγ* gene expression in mature 3T3-L1 adipocytes, suggesting a possible negative feedback loop [39]. Our findings are contrasted by those of Xiang et al. [40], who studied regulation of *Angptl4* by various fatty acids in hepatocytes isolated from the fish *Larimichthys crocea*. These authors showed that 12h treatment with 200μM PA and oleic acid (OA) increased *Angptl4* gene expression, while 200μM ALA, EPA, and DHA did not. Our findings do not support that PA can increase *Angptl4* gene expression; however, difference in species (fish vs mouse), target tissue (liver vs adipose), and treatment duration (12h vs 48h) could explain this discrepancy. Indeed, it is possible that N-3 PUFA require longer treatment durations to influence *Angptl4* gene expression, or that higher dosages of PA are necessary to elicit an increase in *Angptl4* gene expression in 3T3-L1 adipocytes. Regardless, Xiang et al. [40] identified PPARγ agonism as the primary mechanism by which PA increases *Angptl4* gene expression, which aligns with our proposed mechanism, albeit with a different fatty acid species. Lastly, the potent PPARγ ligand PIO demonstrated extremely similar results to DHA across all experiments, thus providing additional support that DHA works through PPARγ to increase *Angptl4* gene expression. These results align with previous work showing that rosiglitazone also increased *Angptl4* expression in differentiated 3T3-L1 adipocytes [41]. Lastly, our study did not elucidate whether DHA or an oxidized metabolite of DHA was mediating these effects, as oxidized DHA metabolites have also been shown to activate PPARγ [42].

Our findings also reinforce the inverse relationship between ANGPTL4 and LPL activity in adipocytes. Specifically, cells treated with *Angptl4* siRNA had significantly higher LPL activity than cells treated with the scramble siRNA. These findings align with a previous study employing *Angptl4* siRNA in 3T3-L1 adipocytes [31]. When *Angptl4* was silenced, both DHA and PIO showed a reduced ability to inhibit LPL activity relative to the same treatments in the scramble siRNA group. While we hypothesized that silencing *Angptl4* expression would completely ablate the ability of DHA and PIO to inhibit LPL activity, our results suggest that the *Angptl4* siRNA was unable to fully negate DHA- and PIO-induced increases in *Angptl4* gene expression. We attribute this again to the fact that siRNA treatment does not achieve a full knock-out but rather a ∼70% reduction in *Angptl4* expression. As such, future studies in isolated primary adipocytes from an ANGPTL4 knock-out mouse would prove invaluable to address this limitation. Additionally, we did not include analysis of LPL activity in *Pparγ* siRNA-treated cells because silencing *Pparγ* reduces *Lpl* expression, thereby confounding any measures of LPL activity in these cells. A possible limitation with measuring LPL activity in whole cell lysates is that this could introduce the possibility that intracellular lipases such as HSL and ATGL may have influenced our LPL activity assay. However, DHA and PIO increased *Atgl* expression, which is counter to the observed reductions in LPL activity. While DHA did reduce *Hsl* expression, PIO did not. This suggests that the change in *Hsl* expression is unlikely to influence our interpretation given the similar impact on LPL activity seen with DHA and PIO. Importantly, we also conducted an activity analysis on heparin-releasable LPL and found similar results to our whole-cell data, albeit with a lower signal and a smaller effect size. While we cannot say for certain why the heparin-releasable LPL activity is lower than the whole-cell activity, it is possible that our culture conditions, namely the absence of insulin from the cell culture media during fatty acid treatments, could reduce the LPL translocation to the cell membrane thus resulting in higher activity in cell lysates relative to membrane-bound LPL. Regardless, this ultimately underscores the importance of continuing this research *in-vivo*; as measures of plasma LPL activity or measuring lipid uptake into tissues using labelled lipid emulsions will provide additional insights. Finally, it remains unclear how well measures of LPL activity in whole cell extracts translate to *in-vivo* LPL activity measurements, namely heparinized plasma or WAT. We therefore believe that our results using both methods to assess LPL activity provide compelling evidence that regulation of *Angptl4* expression is at least partially responsible for the observed inhibition of LPL activity induced by DHA.

Overall, this study provides new insight regarding the mechanism by which DHA can regulate adipocyte LPL activity and, consequently, whole-body lipid homeostasis. Specifically, DHA significantly increased *Angptl4* gene expression and reduced cellular LPL activity. While the effects of DHA on *Angptl4* expression do not appear to be mediated via insulin signaling in adipocytes, we found that DHA regulation of *Angptl4* expression is dependent on PPARγ. The findings of the present study thus provide a strong foundation for future studies aimed at better understanding the relationship between N-3 PUFA and ANGPTL proteins in WAT, and the wider role of N-3 PUFA regulation of whole-body lipid metabolism.

## Author Contributions

PVM and DMM contributed to study conceptualization and design. PVM conducted all experiments. AR conducted the mouse feeding study and tissue collection. LHB and RE contributed to methods development. PVM and DMM performed data analysis. PVM and DMM wrote and edited the manuscript.

