## Supplemental File for "DHA Increases *Angptl4* Gene Expression and Reduces LPL Activity in a PPARγ-dependent manner in Adipocytes"

Table S1 – RT-qPCR Primer Sequences

| Primer | Sequence |
|---|---|
| *Angptl4 - F* | 5’ GGG ACC TTA ACT GTG CCA AG 3’ |
| *Angptl4 - R* | 5’ GAA TGG CTA CAG GTA CCA AAC C 3’ |
| *Nono – F* | 5’ CCC CAC CAA TAC CTG CAA 3’ |
| *Nono - R* | 5’ TTC AGG TCA ATA GTC AAG CCT TC 3’ |
| *Lpl – F* | 5’ ACT CGC TCT CAG ATG CCC TA 3’ |
| *Lpl – R* | 5’ GTT GTG TTG CTT GCC ATC C 3’ |
| *Atgl – F* | 5’ TGA CCA TCT GCC TTC CAG A 3’ |
| *Atgl – R* | 5’ TGT AGG TGG CGC AAG ACA 3’ |
| *Hsl – F* | 5’ CAC AAA GGC TGC TTC TAC GG 3’ |
| *Hsl – R* | 5’ GGA GAG AGT CTG CAG GAA CG 3’ |
| *Pparγ – F* | 5’ TGC TGT TAT GGG TGA AAC TCT G 3’ |
| *Pparγ - R* | 5’ CTG TGT CAA CCA TGG TAA TTT CTT 3’ |
